# Robust sub-network fingerprints of brief signals in the MEG functional connectome for single-patient classification

**DOI:** 10.1101/2024.06.23.599587

**Authors:** Vasiles Balabanis, Jiaxiang Zhang, Xianghua Xie, Su Yang

## Abstract

Recent studies have shown that the Magnetoencephalography (MEG) functional connectome is person-differentiable in a same-day recording with as little as 20 latent components, showing variability across synchrony measures and spectral bands. Here, we succeed with 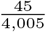 components of the functional connectome on a multi-day dataset of 43 subjects and link it to related clinical applications. By optimizing sub-networks of 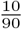 regions with 30 seconds of broadband signal, we find robust fingerprinting performance, showing several patterns of region re-occurrence. From a search space of 5.72 trillion, we find 46,071 of many more acceptable solutions, with minimal duplicates found in our optimization. Finally, we show that each of these sub-networks can identify 30 Parkinson’s patient sub-networks from 30 healthy subjects with a mean F1 score of 0.716 ± 0.090SD. MEG fingerprints have previously been shown on multiple occasions to hold patterns on the rating scales of progressive neurodegenerative diseases using much coarser features. Furthermore, these sub-networks may similarly be useful for identifying patterns across characteristics for age, genetics, and cognition.

## 1 Introduction

Exploring underlying computational relationships in the brain has been a significant area of study in areas such as brain cognition, disease understanding, and clinical diagnostics. The use of electrophysiological signals of the brain, such as from Magnetoencephalography (MEG) scans, provides particularly good temporal resolution and has been found to demonstrate different aspects of disease biomarkers [1] and brain communication architecture [2]. This study explores how person re-identifiable sub-network patterns in the brain can be derived from resting-state MEG in a granular and deterministic way. These granular sub-network patterns are also found to each differentiate between healthy subjects and Parkinson’s patients.

Person re-identification from brain fingerprints has been studied extensively, particularly across Electroencephalography (EEG) [3] and functional-magnetic-resonance-imaging (fMRI) [4]. Studies in EEG finger-printing primarily focus on biometrics applications [5] via machine learning with signal processing methods. In contrast, advances in fMRI fingerprinting have focused on the functional connectome [4] and helped the understanding of several areas of neuroscience, such as cognition [6]. Other modalities of the brain for fingerprinting include MEG [7], structural-magneticresonance-imaging (sMRI) [8], and functional-nearinfrared-spectroscopy (FNIRS) [9]. The prominence of studies and the signal nature of EEG and fMRI make it feasible for methods in signal processing and the functional connectome for these modalities to be applicable in MEG.

Existing studies for MEG fingerprinting explore several computational approaches, primarily on the functional connectome to do person re-identification. Studies performing static connectivity fingerprinting on the functional connectome include amplitude-based connectivity [7, 10, 11, 12, 13] and phase-based connectivity [10, 11, 12, 14, 15, 13, 16, 17]. In [7, 18, 12], connectivity is done on sensor-space data. Dynamic connectivity fingerprinting on the functional connectome includes signal avalanching [19] and spectral-coupling [20]. Approaches without connectivity include spectral activity [7, 11, 12, 18, 21] and temporal activity [18, 21, 22].

In Silva Castanheira et al. [7], the uniqueness of the MEG functional connectome fingerprints for 47 subjects was found to be robust across two sessions with an average duration of 201.7 days. In same-day data, these fingerprints have shown increased re-identifiability patterns in cognitive tasks [11]. It shows reduced patterns in levels of sleep deprivation [14] and progressive neurodegenerative diseases such as Parkinson’s [15], Mild Cognitive Impairment (MCI) [13], Multiple Sclerosis (MS) [16], and Amyotrophic Lateral Sclerosis (ALS) [17]. The fingerprint has shown to have a statistically significant explainable variance in predicting the Unified PD Rating Scale part III (UPDRS-III) [23] for Parkinson’s disease and rating scales for MCI, MS, and ALS.

The overall fingerprinting robustness and granularity of the MEG functional connectome remain partially unexplored in literature. For granularity, instances of functional connectome feature-reduction currently include [12, 13, 15, 16, 17, 20, 19]. In Haakana et al. [12], it was found that only 20 latent components were needed for person re-identification for same-day data. In Dimitriadis et al. [20], it was shown that only 76 dynamic spectral-coupling components were needed for person re-identification for multi-day data. In Sorrentino et al. [19], subsets of regions representing dynamic signal avalanche components were needed for person reidentification for multi-day data.

In progressive neurodegenerative studies [15, 13, 16, 17], it was found that by only selecting a threshold of values of the functional connectome consistent in the diseased groups, the patients’ re-identifiability and rating scale prediction was improved.

To address these patterns in literature, an effective solution to perform feature reduction while preserving fingerprinting performance, such as meta-heuristic optimization algorithms, can be used. Existing EEG biometrics studies have previously used optimization algorithms with temporal and spectral features to find salient electrodes for fingerprinting [24, 25]. A combinatorial optimization approach has not to date been applied to the selection of regions in the MEG functional connectome for person re-identification capabilities.

In this study, we used meta-heuristic optimization to perform fingerprinting on sub-sampled features in the MEG functional connectome. We did feature reduction by taking sub-sets/sub-networks of regions in the Automated-Anatomical-Labeling (AAL) atlas. These sub-networks were optimized with fingerprint metrics across a search space of 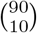 or 5.72 trillion possible sub-networks. We obtained 46,071 maximally optimal sub-networks with varied patterns, with only 263 sub-network duplicates between 24,310 independent runs. With these optimal sub-networks we acquired a granular representation of the salient regions in the functional connectome while also providing a significant increase in feature dimensionality. Finally, using this increased dimensionality, we observed non-linear patterns across these subnetworks that differentiated 30 Parkinson’s disease patients and 30 healthy subjects with an accuracy mean of 0.740 ± 0.004SD and an F1 score mean of 0.716 ± 0.090SD.

## 2 Results

Our findings show 100% re-identification performance of 46,071 sub-networks from 10 regions in the AAL atlas across 43 subjects. The search space was 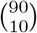, which is 5.72 trillion sub-networks, demonstrating a large span for possible optimal solutions. Our use of multi-day data (mean: 6.07 days, ± 6.14SD) ranging from 1-27 days shows the robustness of the re-identifiability patterns in these sub-networks. We further observed graph measures relating to the performance and occurrence of regions in the sub-networks that provide insight into the region salience and co-occurrence structure of the search space.

Using our sub-networks on an external dataset of 30 Parkinson’s patients and 30 healthy subjects, we find that the non-linear patterns across 40,000 of the subnetworks demonstrate the differentiation of Parkinson’s patients from healthy subjects. The F1 score test performance of sub-networks on all subjects was mean: 0.716 ± 0.090SD. In contrast, using the null distribution, the F1 score test performance was 0.550 ± 0.136SD.

### 2.1 Finding sub-networks

We first structurally mapped and source-localized MEG signals of 43 subjects using Linearly Constrained Minimum Variance (LCMV) beam-forming (see Methods 5.5). We then averaged across all voxels within regions in the AAL atlas. We used only the first 90 regions of 116 for analysis. Each region contained a 115 s signal in broadband frequency (0.1-150 Hz). We then took the functional connectome by calculating the Pearson correlation between all possible pairs of 90 regions for 30 s of signal, resulting in a 90 × 90 correlation matrix (see Methods 5.6).

To perform fingerprinting, we then once more used the Pearson correlation of the upper triangle of the subject’s first session with that of their second session. We compared whether the subject’s cross-session identifiability score was higher than the correlation of the subject’s first session with all the other sessions of the other subjects. The session that returned the maximum correlation score gave us the re-identification prediction.

Furthermore, we calculated a score known as ‘differential identifiability’ originally used in [26]. This gave a representation of the margin between intra and inter-identifiability across all subjects (see Methods 5.7).

To start the analysis, we aimed to find re-identifiable patterns in sub-networks/sub-sets of regions in the MEG functional connectome. A null distribution size of 10,000 random samplings of various lengths was taken until a threshold was found where a significant percentage (95%) of randomly selected sub-networks resulted in significant accuracy (+95% or 41/43 successful classifications). This condition was met when using at least 36 regions.

Next, we decided to narrow down a region number that we could use for optimization. We took the null distribution for the number of regions in which only the top 5% of their distribution represented 95% or more accuracy. This was to ensure that later optimized subnetworks were statistically better at fingerprinting than random. From 10,000 null distribution samplings, suitable candidates for this analysis ranged from 8 to 12 regions. We decided on 10 regions based on the poor performance of 8 and 9 regions in later optimization steps. 10 regions maintained the stated distribution of accuracy requirements (see Supplementary Fig. 1).

We used the Simulated Annealing optimization algorithm (see Methods 5.8), which iteratively randomly swapped regions in the sub-network to converge to an optimal fitness score. The fitness function was based on recognition accuracy and differential identifiability (see Methods 5.8). After fine-tuning our optimization configuration locally, we would run it several times using parallel computing on the supercomputer available to our institute.

After optimization, which would be validated on evaluations of two sets of two 30 s segments of the MEG signal from different sessions, we would filter out solutions that did not have 100% (43/43) accuracy on both instances of cross-session fingerprinting. We would later filter solutions that did not have 100% cross-session fingerprinting accuracy on a third unseen set of two 30 s segments. For later analysis, we additionally independently sampled only the best solution from each run of Simulated Annealing to avoid sub-network region and performance similarities between successive iterations of the algorithm.

Next, we ran our optimization using a supercomputer. We first decided to perform an exploratory optimization search to choose our data hyperparameters. In a study by Fraschini et al. [27], it was found that as little as 10 s of signals show consistent graph connectivity patterns in EEG. Additionally, studies have found 30 s of the MEG functional connectome to be stable [28] and robust [7] for fingerprinting. In [7], there are also indications that a larger signal duration can boost fingerprinting performance. Furthermore, we found performance variations in the number of regions from our initial optimization searches that were run locally.

We, therefore, grid-searched the duration of the MEG signal (10 s, 30 s, 50 s) and the number of regions used (10, 20, 30, 40). Differential identifiability was used to evaluate performance. We ran 5,000 optimization runs for each of these 12 conditions, where their convergence is shown in Fig. 2.

**Figure 1:**
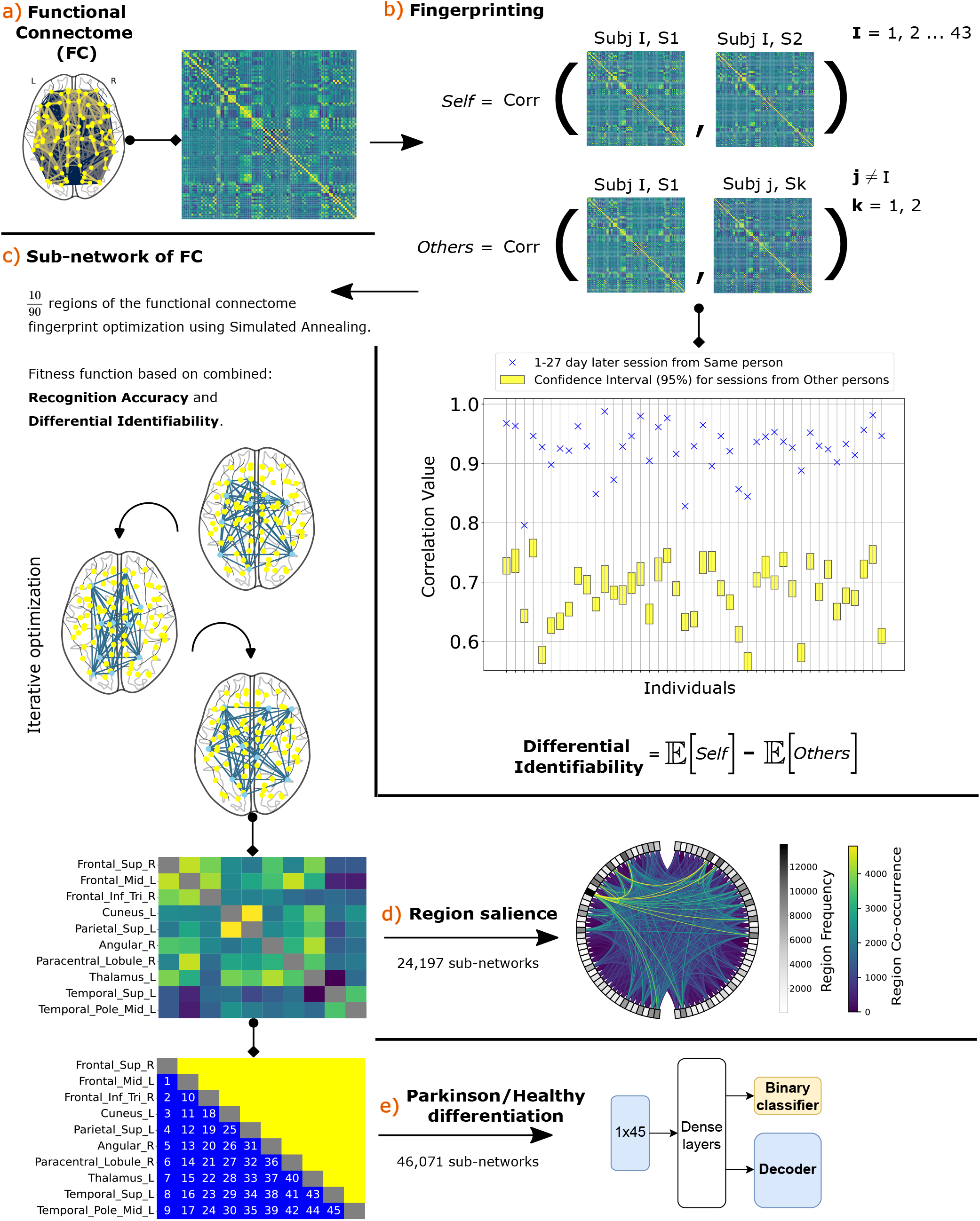
Analysis pipeline. **a)** Pearson correlation is calculated between all regions’ signals, creating a 90×90 correlation matrix (see Methods 5.6). **b)** To perform fingerprinting, Pearson correlation is calculated between correlation matrices to generate a confidence score for the person it belongs to. The highest score returned from 85 sessions, one of which belongs to the cross-session of the same subject, indicates the prediction (see Methods 5.7). **c)** Simulated Annealing optimization function (see Methods 5.8) with a fitness function of combined differential identifiability and recognition accuracy is used to generate optimal sub-networks of 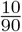 regions in the AAL atlas. **d)** These sub-networks were combined into a larger connected graph based on co-occurrence and analyzed (see Methods 5.9) in terms of frequency of occurrence, co-occurrence frequency, and **not shown in the figure:** assortativity, local region neighborhood clustering, and performance of subject-specific region co-occurrence. **e)** These sub-networks were fed into a dual-objective stacked auto-encoder model (see Methods 5.10), which differentiated between the sub-networks of 30 Parkinson’s and 30 healthy control subjects.

**Figure 2:**
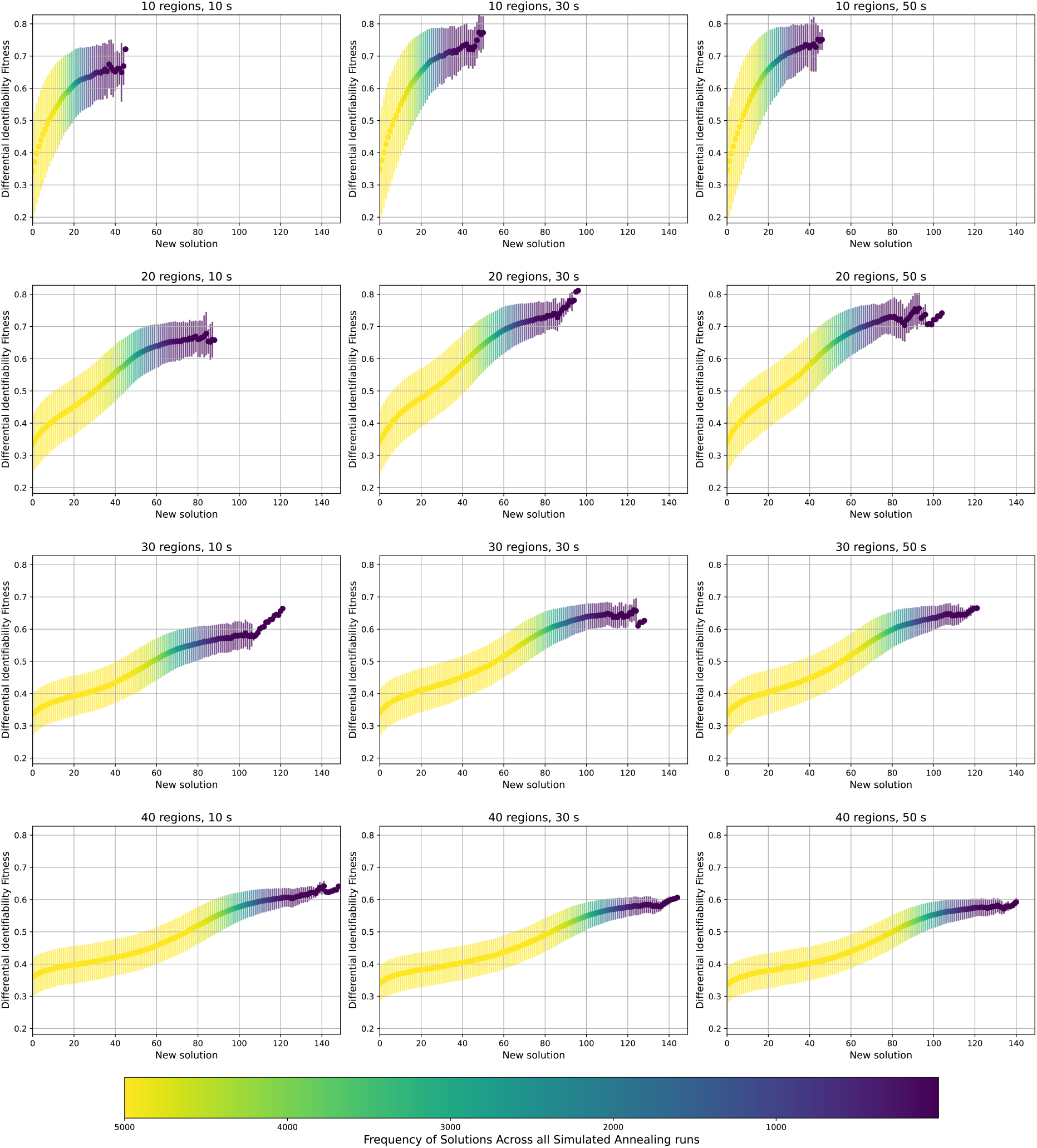
Differential identifiability fitness convergence of 5,000 Simulated Annealing runs for each of 12 conditions. The conditions are MEG signal duration and number of regions in the sub-network. This shows the standard deviation (vertical spread), mean (central line plot), and frequency (hue) of runs for 5,000 runs of each condition reaching a certain performance. Conditions are across the number of regions and duration in the MEG signal.

We finally made the decision to use 30 s as the primary duration to explore based on our affirming grid-search results and the results of the MEG resting-state connectivity stability study [28] and robust cross-session fingerprinting performance by Silva Castanheira et al. [7].

Following this, we ran a total of 100,000 optimizations of Simulated Annealing on 30 s using 10 regions in the MEG functional connectome of the AAL atlas. After acquiring results and filtering optimal subnetworks, we obtained 48,327 sub-networks, which after de-duplication were 46,071. When we independently sampled from runs, we obtained 24,310 sub-networks, which were 24,197 after de-duplication. Between all independent runs for 48,327 solutions, there were 263 duplicate sub-networks. The differential identifiability score for 24,197 sub-networks was mean: 0.649 ± 0.127SD, composed of the intra-identifiability (mean: 0.918 ± 0.013SD) and inter-identifiability (mean: 0.270 ± 0.131SD), shown in Fig. 3.

**Figure 3:**
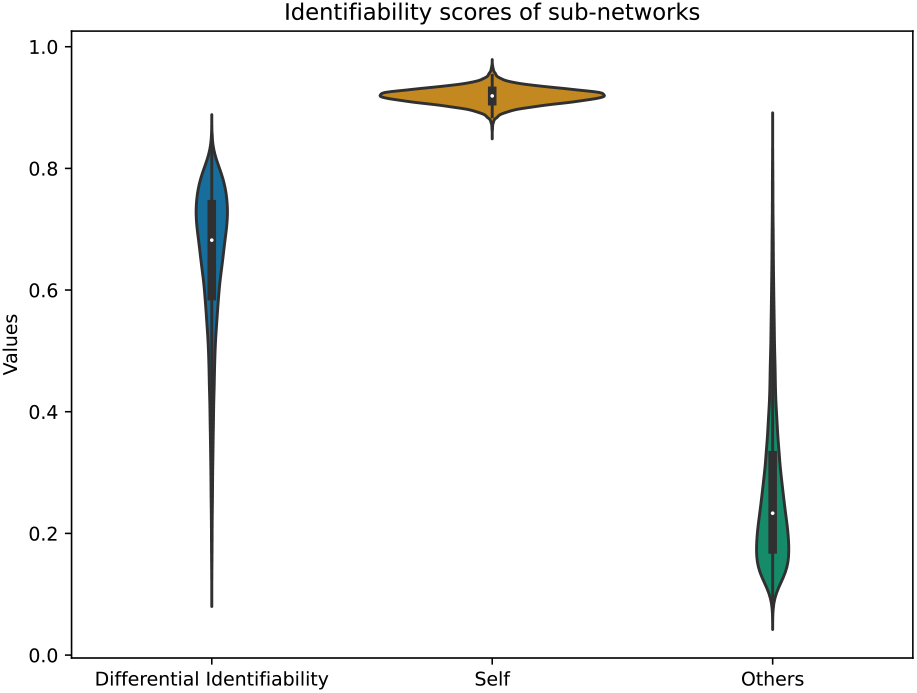
Sub-network identifiability scores. This is based on Equations in Methods 5.7. These were de-duplicated and independently sampled and display scores from 24,197 sub-networks.

### 2.2 Investigating sub-networks

Once our 24,197 sub-networks were obtained, we constructed a graph where edges represented the frequency of co-occurrence between regions. We took graph measures to observe any patterns and structures present in the optimized search space.

We first explored the frequency and co-occurrence in which regions appeared in these sub-networks, as shown in Fig. 4. We further observed the average co-occurring differential identifiability with the top 10% scores (see Supplementary Fig. 2).

**Figure 4:**
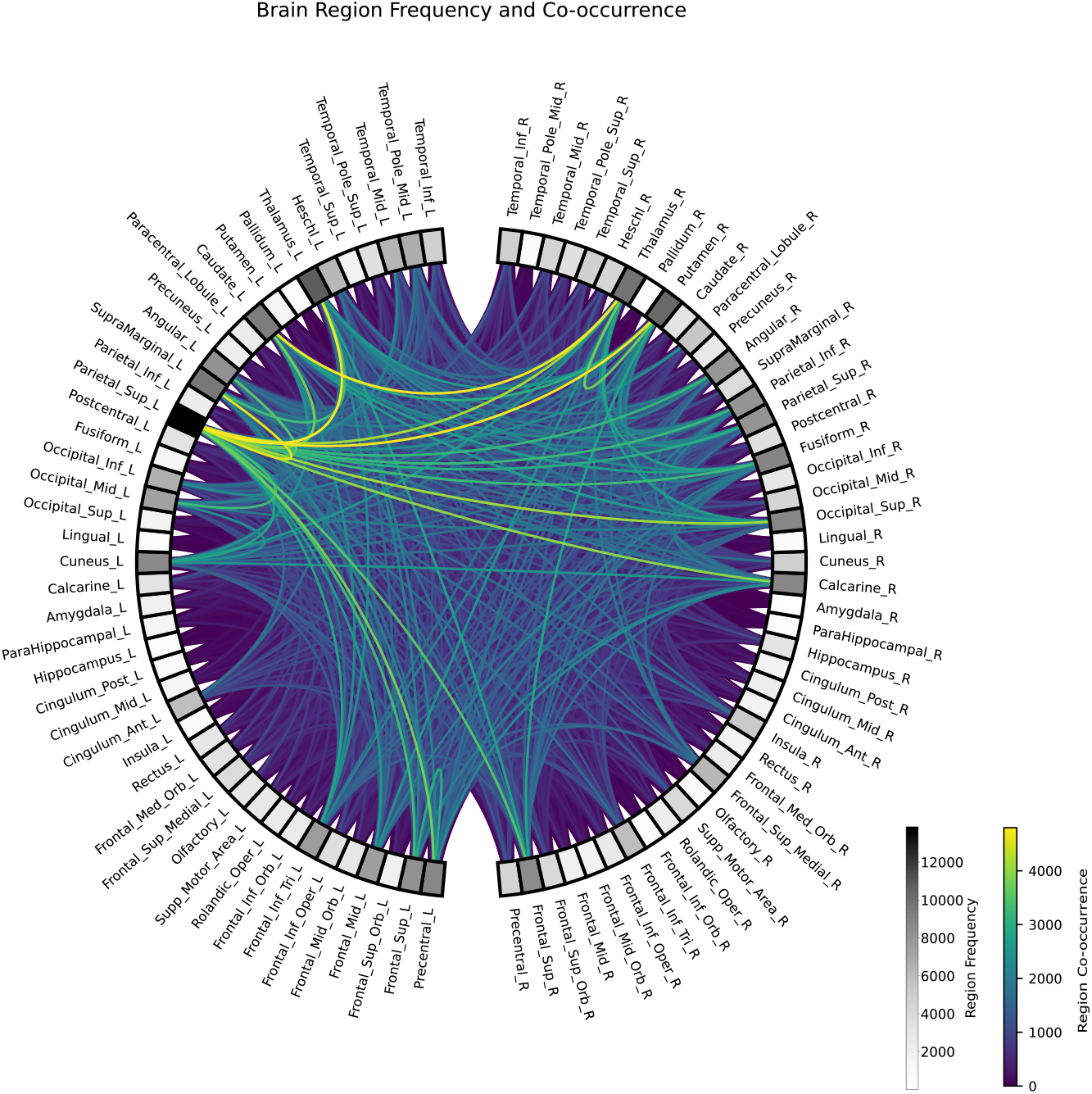
Region co-occurrence and frequency across 24,197 filtered sub-networks. Each of the sub-networks is from independently sampled Simulated Annealing runs.

We then explored the graph structure of the co-occurrences of regions in the 24,197 sub-networks. We first observed **1) assortativity** (see Methods 5.9.1), representing the tendency of more frequently occurring regions to co-occur with less frequently occurring. The score was found to be 0.002. The assortativity was also taken in with the absence of more frequently occurring regions, shown in Supplementary Fig. 3.

Second, we observed the **2) local region neighborhood clustering** (see Methods 5.9.2) to see whether a region’s community of co-occurrences have at least one co-occurrence with each other in the 24,197 sub-networks. This is shown for each region against its frequency of occurrence and its total number of co-occurring regions in its neighborhood (see Supplementary Fig. 4). This gave a representation of region-specific co-occurrence and the tendency for communities of regions to co-occur together. The clustering coefficient scores’ mean was 0.964 ± 0.013SD (0.943-1.000).

Lastly, we measured **3) region co-occurrence by subject-specific performance** (see Methods 5.9.3). This was done by observing how many times a region was maximally beneficial to the performance of other regions when co-occurring in the 24,197 sub-networks. This was done 43 times for each subject to observe heterogeneity. Performance was based on the average intra-identifiability of their co-occurrence. We used intra-identifiability instead of differential identifiability to remove the bias that is included from other subjects (see Equation 5) and thus give a more universal score outside of our cohort.

The subject-specific intra-identifiability scores averaged between all sub-networks showed a mean of 0.919 ± 0.040SD (0.798-0.981), showing the level of heterogeneity in performance for each subject. The degrees of regions were taken by how many times the region was with maximal performance to others. The degree divergence (see Equation 11) across subjects was found to be mean: 3.813 ± 1.486SD (2.405-10.173), with the degree mean as: 1.675 ± 0.064SD (1.533-1.889).

Furthermore, we observed the number of regions that were not beneficial to any others. This was the leaf number, and the mean was 63.63 ± 3.63SD (58-76) from a maximum value of 89 regions. These results show that particular regions in each subject-specific graph contributed to the maximal score.

In Supplementary Fig. 5, the tendency for which of the same regions is found to have maximum co-occurrence performance in all subjects.

### 2.3 Application of sub-networks

Motivated by previous studies that showed variations in fingerprinting scores between diseased and healthy groups [15, 13, 16, 17], we aimed to observe any between-group patterns of our sub-networks in Parkinson’s disease and healthy subjects. In [15], it was found that Parkinson’s patients had reduced re-identifiability patterns in the beta band (13-30 Hz) from phase linearity measurement (PLM) connectivity [29] on 210 s. These beta band patterns were also shown to be a statistically significant explainable variance in predicting the UPDRS-III rating scale [15].

We tested these sub-networks on the single-session MEG recordings of 30 Parkinson’s patients and 30 healthy subjects from the OMEGA dataset (see Supplementary Table 2). We used this dataset despite age differences between healthy subjects in the OMEGA data (mean: 67.11 ± 8.35SD) and healthy subjects in our data (mean: 22.45 ± 3.06SD). From the 46,071 de-duplicated unique sub-networks that we found in our dataset, we obtained the functional connectome matrices of these sub-networks for Parkinson’s patients and healthy subjects. To begin, we sampled functional connectomes from only one 30 s segment from each of the 46,071 sub-networks for all subjects in the OMEGA dataset. This provided for each sub-network sample a 10 × 10 correlation matrix, which consisted of a 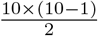 : 45-size feature vector of unique region-pair values. We used each of these sub-networks as samples for a deep-learning model that tried to differentiate the diseased sub-networks from the healthy sub-networks. We decided on a lightweight dual-objective stacked auto-encoder deep learning model for binary classification (see Supplementary Methods 2). We tested with 6-fold cross-validation, testing on each fold an equal number of 5 healthy subjects and 5 diseased patients for 100 epochs (see Methods 5.10).

Our results show that Parkinson’s disease patients’ sub-networks are differentiated from healthy using 40,000 of the 46,071 identified optimal sub-networks. This was with a mean accuracy of 0.740 ± 0.004SD compared to 0.605 ± 0.002SD from the null distribution of 40,000 sub-networks.

There were significant accuracy performance differences between the number of features (1,000, 2,000, 5,000, 10,000, 20,000, 40,000) in the null distribution sub-networks and optimal fingerprint sub-networks. This is shown in Fig. 5.

**Figure 5:**
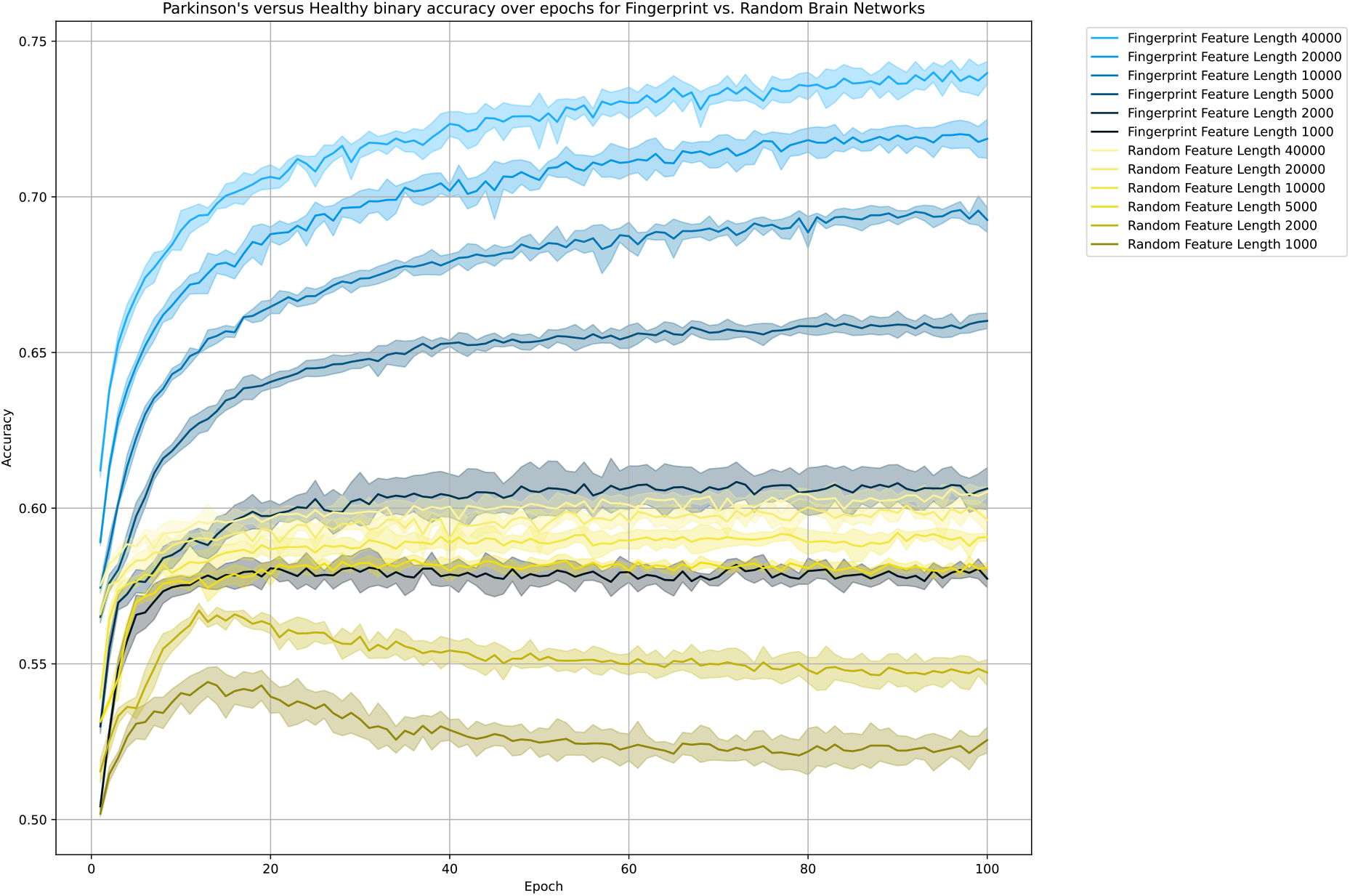
Test accuracy across epochs on the dual-objective stacked auto-encoder. This is elaborated in Methods 5.10. The vertical margins are standard deviations based on the spread of folds in 6-fold cross-validation, while the line plots represent the mean.

The F1 score (see Supplementary Methods 3) of each of the 40,000 optimal fingerprint sub-networks is shown in Fig. 6, with a mean F1 score of 0.550 ± 0.136SD for the null distribution and 0.716 ± 0.090SD for the optimal sub-networks.

**Figure 6:**
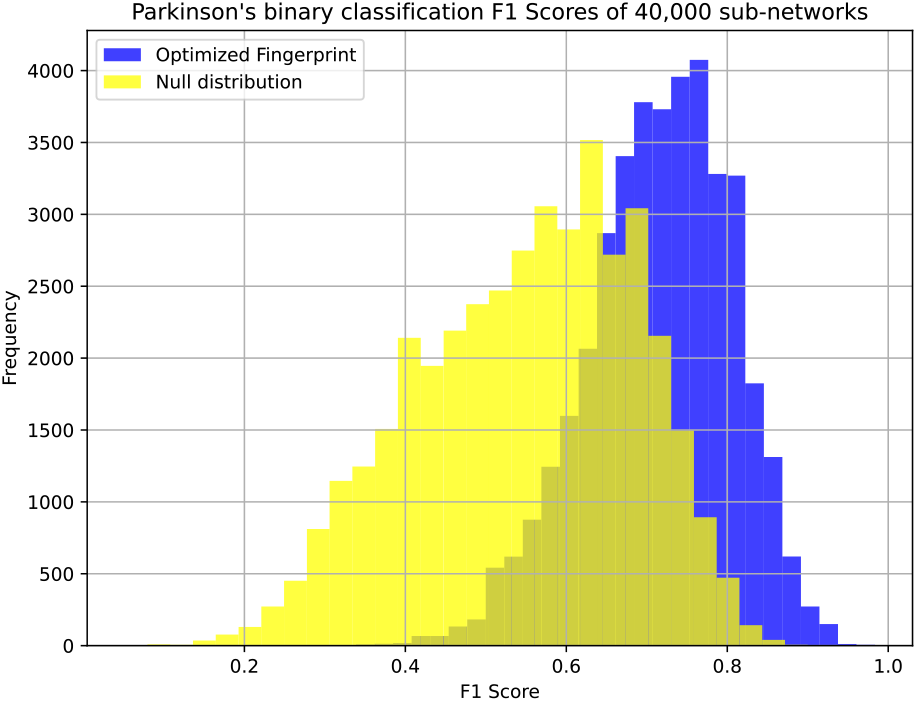
F1 score histogram across 40,000 sub-networks. This is across all cross-validation test samples for Parkinson’s classification using dual-objective stacked auto-encoder (See Supplementary Methods 2).

The consistency of correct classifications for sub-networks across individuals is shown in Supplementary Fig. 6.

## 3 Discussion

Our findings demonstrate that only 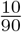 regions from the MEG functional connectome are needed to perform fingerprinting. We find 46,071 of the sub-network fingerprints to be consistent with 100% person reidentification performance of all of our 43 subjects, validated on three sequential segments of signal. These sub-networks were found from a possible search space of 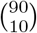 or 5.72 trillion possible sub-networks.

We further explored the regions present in these sub-networks and found tendencies in which regions co-occurred, representing the patterns and structure derived from the optimized search space.

Lastly, we used these sub-networks and found related applications for Parkinson’s disease. This is shown by our binary classification performance of mean: 0.716 ± 0.090SD F1 score across 40,000 sub-networks between 30 Parkinson’s patients and 30 healthy subjects. This demonstrates the underlying sub-network fingerprint patterns of the MEG functional connectome that relate to the disease. This is using only 40,000 of the 5.72 trillion possible sub-networks. The relevance of our MEG sub-network fingerprint to progressive degenerative diseases other than Parkinson’s is shown by the following studies [13, 16, 17], which found their rating scale is related to significantly less granular features than we have used.

### 3.1 Re-identifiability of subjects

Our method demonstrates 100% person re-identification performance using sub-networks with deterministic regions/features. The use of multi-day data (1-27 days) for robustness, only done in MEG fingerprinting studies by [7, 20, 19, 18] further strengthens the significance of these sub-networks. Our findings show that 10 out of 90 regions are feasible for our method from a search space of 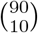, where each sample represents 45 components.

In Haakana et al. [12], 20 latent components were found to have 90%+ fingerprinting accuracy on the functional connectome for same-day data. This was found using phase locking value (PLV) from 360 s of signal duration. The performance was reduced to 80% when tested on unseen subjects. Our sub-networks should similarly be investigated on a subject group external to our optimization to verify the universality of our fingerprints. Haakana et al. also used different spectral band-widths and synchrony measures, showing significant variations in intra-identifiability and inter-identifiability scores across them. Our study presents a possible exploration into the characteristics of these approaches for our sub-network fingerprints.

Our Simulated Annealing results showed performance variations between the number of regions (10, 20, 30, 40) and MEG signal duration (10 s, 30 s, 50 s) as shown in Fig. 2. Our findings found strongest differential identifiability convergence for (10 regions, 30 s), (10 regions, 50 s), (20 regions, 30 s), and (20 regions, 50 s). The higher differential identifiability performances in a lower number of regions may be attributed to the smaller search space, making it easier to converge to an optimal solution. An additional advantage to using fewer regions was that it provided stronger salience of regions and dimensionality reduction useful for later analysis.

The choice of MEG signal duration aside from what we have explored (10 s, 30 s, 50 s) should be further investigated. Though it has been indicated that 30 s of the signal contains stable, functional connectome connectivity patterns in MEG [28], in fMRI, the functional connectome’s connectivity fingerprint performance has been shown to vary with signal duration [30] across temporal scales. Furthermore, in MEG, it appears that only 1 s of temporal features from the functional connectome may be required for person re-identification using deep learning [22]. The precise range of temporal duration for fingerprinting capabilities in literature is unexplored. Further investigation into the robustness across signal duration may be useful for future fingerprint analysis.

Lastly, our study involves a 1-27 day interval and shows 95%+ re-identification accuracy for 36-90 regions of 10,000 randomized sub-networks in the functional connectome. In contrast, Silva Castanheira et al. [7] with 30 s of broadband (0-150 Hz) used 47 subjects and a 201.7 day average between sessions. They found 83.8% re-identification accuracy from the full-brain functional connectome. Additionally, Silva Castanheira et al. found no linear relationship between the fingerprinting performance of subjects and the duration of their sessions. Further investigation should be explored using our upscaled features to find patterns that may better relate to the number of days between sessions.

### 3.2 Investigation of salient regions

Investigation into the different numbers of regions (10, 20, 30, 40) that we used and signal duration (10 s, 30 s, 50 s) demonstrated across all conditions similar region frequency and co-occurrence frequency patterns. These patterns are shown for 10 regions with 30 s in Fig. 4.

We found 263 optimal sub-network duplicates between 24,310 independent runs, demonstrating a large remaining optimal landscape for solutions.

The assortativity score of 0.002 demonstrated that regions of the 24,197 sub-networks did not co-occur based on regions with many to many (maximum 1) or less to many (minimum −1) frequency of occurrence. This demonstrates a neutral tendency for regions to co-occur in terms of region frequency of occurrence. This further indicates a more heterogeneous and potentially larger portion of the overall search space of 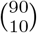 that may be explorable.

Next, by observing the local region neighborhood clustering, we could see whether regions co-occurred at least once in region-specific communities. The 0.964 ± 0.013SD (0.943-1.000) score showed that region-specific co-occurring regions were mostly co-occurring with each other at least. This shows the diversity of region co-occurrences in the search space across all regions.

The region clustering showed the tendency for regions with fewer regions co-occurring at least once to have a slightly higher clustering coefficient value. It also generally showed that these same regions had a lower frequency of occurrence. See Supplementary Fig. 4 to see these observations in our 24,197 sub-networks. These patterns, though representative of a limited portion of the landscape, could hypothetically be explained. It is possible that some regions have more homogeneous neighborhoods, while others have more heterogeneous. This may indicate that regions with more homogeneous communities have a comparatively smaller set of total optimal solutions from the 5.72 trillion search space, aside from the obvious indication of less frequency of occurrence. It may also indicate that particular region co-occurrences are generally more suitable for the optimality of specific regions.

An extension to this study could explore fine-tuning an optimization algorithm that finds solutions limited to particular regions. This could further investigate the co-occurrence landscape of regions in the search space. Furthermore, a more in-depth investigation into how communities cluster based on the frequency of co-occurrences should be considered for a detailed view.

Finally, we observed how many times each region had the maximal value of average intra-identifiability score when co-occurring with other regions. Since we took the subject-specific intra-identifiability on each of the 24,197 sub-networks, we were able to observe heterogeneity across all subjects.

In our initial optimization search, from the 24,197 sub-networks, as found in Section 2.1, there was a mean: 0.919 ± 0.013SD (0.852-0.975) score for the intra-identifiability across all sub-networks. When averaging across all sub-networks for each subject, we instead observed a mean of 0.919 ± 0.040SD (0.798-0.981). The variation between sub-network scores and subject-specific scores shows the span of uniqueness of the subject-specific intra-identifiability.

We additionally observed degree divergence and leaf number. These measures described the distribution of regions with maximal performance co-occurrence, where a higher value of degree divergence indicates the broadness for regions to be maximal in performance to other regions (see Equation 11). A higher leaf number indicates more regions that are not maximal in co-occurrence performance to other regions.

The degree divergence distribution had a mean of 3.813 ± 1.486SD (2.405-10.173), and the leaf number was 63.63 ± 3.63SD (58-76). Supplementary Fig. 5 shows that many of these degrees seem to come from similar regions across all subjects. These results show that particular regions are responsible for the maximal co-occurrence performance in all 43 subject-specific graphs. Further investigation should numerically examine the margin between the maximal regions and non-maximal regions. Exploring this would give a detailed view of co-occurring intra-identifiability performance. The heterogeneity of fingerprint values may be attributed further to demographic data such as the age, the handedness[7], or the cognitive scores [10] of the subject.

In future studies, all of our sub-network region-salience findings should be linked with the network groups in Yeo’s 7 functional networks [31], which were found to contribute heterogeneously to fingerprinting in [7, 11]. Additionally, the region salience in our sub-networks should further be investigated for measuring the fingerprinting across demographics [7], cognitive characteristics [11] and genetic similarities [32].

### 3.3 Clinical application of these fingerprints

Based on findings by [15, 16, 13, 17], we aimed to use our 46,071 discovered sub-networks to see if they carry information sensitive for identifying patients with neurodegenerative diseases. Due to the open-access availability of the OMEGA dataset, we were able to use 30 Parkinson’s and 30 healthy subjects with our upscaled features to differentiate sub-network patterns of the disease.

We decided to perform binary classification using deep learning to see if these sub-networks contain any differentiable patterns. Due to fingerprint score variations in [15, 16, 13, 17] between diseased and healthy subjects, we initially explored using the fingerprinting scores as features with various machine-learning methods for binary classification. These attempts were based on using either intra-identifiability or differential identifiability fluctuations between our 46,071 optimal sub-networks segments sampled from two within-session segments in the functional connectome. These attempts were not successful, prompting us to try something else. We instead took unique pair values in the 30 s snapshot of the functional connectome, resulting in a 45-component feature vector.

Our deep learning model followed a basic architecture: dual-objective regression and binary classification (see Supplementary Tables 3, 4, 5). It was purposefully lightweight for reproducibility and simplicity. We trained on a maximum of 40,000 sub-network samples, each with a feature size of 45. Each sub-network had different indices for the same region, which made it impossible to map a dimension to a specific region. This meant that the model was learning the general patterns of the functional connectome sub-networks and, from that alone, differentiating patterns of the disease.

As shown in Figs. 5 and 6, our findings indicate that the sub-network fingerprints identified in the CUBRIC dataset contribute significantly more to the classification of Parkinson’s disease in the OMEGA dataset than random. The distribution of F1 scores in 40,000 sub-networks Fig. 6 shows that 45-component vectors contain information to differentiate the disease between 30 Parkinson’s and 30 healthy. This was done with a mean F1 score of 0.716 ± 0.090SD compared to 0.550 ± 0.136SD from the null distribution. This corroborates prior indications that MEG fingerprint patterns vary according to Parkinson’s disease.

The application of these sub-networks in other areas is numerous. Based on previous studies, this finding is likely to be applicable to the analysis of levels of sleep deprivation [14], different cognitive tasks [11] and genetic characteristics [32]. Cognitive scores have also contributed to statistically significant explainable variance for predicting functional connectome values [10]. These sub-networks are also applicable in progressive neurodegenerative diseases such as Mild Cognitive Impairment [13], Multiple Sclerosis [16], and Amyotrophic Lateral Sclerosis [17].

In these neurodegenerative studies [15, 13, 16, 17], the explainable variance for prediction of rating scales of the diseases from the fingerprint was shown to be statistically significant. They used PLM synchrony measures in specific spectral bands from the functional connectome using 210 s signal duration. They also used sub-sampled values in the functional connectome to achieve their best performance. In our study, we showed that by using broadband Pearson correlation for 30 s, we could achieve disease differentiation. This was with each of 40,000 of our sub-networks with a significant F1 score of 0.716 ± 0.090SD compared to 0.550 ± 0.136SD from the null distribution. This is despite the volume conduction effects that Pearson correlation connectivity is known to have for estimating the functional connectome. Our upscaled features may present the beginning of more effective monitoring and prediction of these diseases. This is given the search space is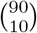, approximately 5.72 trillion possible sub-networks and our discovery of 46,071 sub-networks with only 263 independently observed sub-network duplicates. Spectral bands and synchrony estimations of the functional connectomes can additionally be taken for our 46,071 optimal sub-networks. Further recreating this study on different spectral bands and synchrony measures may uncover deeper characteristics of these diseases.

Further applications include inspirations from fMRI fingerprinting studies. The fMRI functional connectome connectivity fingerprints have shown to differentiate between ages and between genetic similarity [33], have shown reduced re-identifiability patterns in psychedelic drug consumption [34, 35], and have shown to be statistically significant in the prediction of cognitive psychometrics [36]. These could be investigated with MEG, potentially using our sub-networks.

Additionally, the multi-modal relationship of fMRI with MEG for fingerprinting should be further explored with these sub-networks. It has already been explored [10] in the correlation between consistent fingerprint values in the functional connectome for fMRI and MEG across particular regions and the 7 Yeo functional networks.

Extensive investigation is necessary into the nature of the information contained in the sub-networks. This information could potentially be useful for a range of other cognitive and disease-related studies.

## 4 Conclusion

In conclusion, we have shown that by using a minimal number of 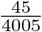 feature values in the MEG functional connectome, a strong, deterministic multi-day fingerprinting performance can be achieved across 46,071 instances. Additionally, using varied samples of these features, we have shown that Parkinson’s disease can be identified from healthy subjects with a mean F1 score of 0.716 ± 0.090SD. This shows major possible applications for the monitoring and prediction of progressive degenerative diseases. This also shows applications for the analysis of general electrophysiological patterns in the brain under different conditions. We hope that future research aiming to understand the characteristics of the fingerprint will benefit from our work.

## 5 Methods

### 5.1 CUBRIC dataset

43 healthy subjects were recruited from a local participant panel consisting of undergraduate and postgraduate students (31 females, age range 18-30 years, mean age: 22.45 ± 3.06SD years). All subjects underwent a T1 MRI session and completed two MEG sessions. No subject reported a history of neurological or psychiatric illness. The study was approved by the Cardiff University School of Psychology Ethics Committee. All subjects gave written informed consent. The use of two cross-day sessions (range 1-27 days, mean days: 6.07 ± 6.14SD) enabled us to observe fingerprint re-identification performance under robust conditions.

### 5.2 Open-access dataset

We used imaging data from the Open MEG Archive (OMEGA) [37], which contains disease and control population-derived cohorts. We chose all 30 healthy subjects between 50 and 83 years old (mean age: 67.11 ± 8.35SD years) and 30 Parkinson’s patients between 48 and 80 years old (mean age: 67.06 ± 8.73SD years). The inclusion of the OMEGA dataset enabled us to incorporate clinical observations into our findings.

### 5.3 MEG and MRI data acquisition for CUBRIC dataset

Whole-head MEG recordings were acquired in a magnetically shielded chamber using a 275-channel CTF radial gradiometer system (CTF Systems, Canada) at a sampling rate of 1,200 Hz. One sensor was turned off during recording due to excessive noise. An additional 29 reference channels were recorded for noise cancellations, and the primary sensors were analyzed as synthetic third-order gradiometers. Continuous horizontal and vertical bipolar electrooculograms (EOG) were recorded to monitor blinks and eye movements. Subjects were seated comfortably in the MEG chair, and their head was supported with a chin rest to minimize head movement. For MEG/MRI co-registration, the head shape with the position of coils was digitized using a Polhemus FAS-TRAK (Colchester, Vermont). Subjects were instructed to rest with their eyes open and fixate on a red dot with a grey background, presented through a back projector. Each recording session lasted approximately 5 minutes. 43 subjects underwent two resting-state MEG sessions on different days (mean: 6.07 ± 6.14SD interval, with a range of 1 to 27 days between two sessions).

Subjects underwent high-resolution T1-weighted magnetization prepared rapid gradient echo scanning (MPRAGE: echo time 3.06 ms; repetition time 2,250 ms, flip angle 9°, the field of view=256 × 256 mm, voxel size 1 × 1 × 1 mm).

### 5.4 MEG preprocessing

MEG data is pre-processed following an analysis. Continuous raw MEG data was imported to Fieldtrip, down-sampled to 300 Hz, and band-passed between 0.1 to 150 Hz. Subsequently, data was notch-filtered at 50 Hz for the CUBRIC dataset and 60 Hz for the OMEGA to remove line noise. Visual, cardiac, and muscle artifacts were removed using ICA decomposition. Identification of visual artifacts was aided by simultaneous EOG recordings where they were available in the CUBRIC dataset. Between two and twelve ICA components were removed for each subject.

### 5.5 MEG source localization

The brain, skull, and scalp boundary surfaces were generated from Fieldtrip using the MRI scans. An automated procedure was used to align these data to the SPM-8 coordinate system [38]. The scalp surface was used to align the structural data with the MEG digitizers. The aligned MEG gradiometers, inner skull surface, and cortical surface were then used to construct a realistic, subject-specific, single-shell forward model. Dipole orientations were fixed normal to the cortical surface. The MEG lead field was created using a volume-conduction model with the boundary element method. Then, MEG activity in atlas-labeled voxels was reconstructed using the LCMV beam-forming source localization algorithm [39].

Head position was estimated from the circumcenter of three head localization coils in each trial. Head movement trajectories containing transformations and rotations were z-transformed and regressed out from source-level MEG time series.

The cortical surface was aligned to the AAL atlas (116 regions) in Fieldtrip. After applying the LCMV weight vector for each voxel to the MEG sensors, the time series of each region of interest (ROI) was calculated as the average of all voxels within the ROI. Only the first 90 ROIs were kept due to the unreliability of source localization in the vermis and cerebellum regions.

### 5.6 MEG-based functional connectome

Functional networks were constructed using raw Pearson correlation within broadband frequency (0.1-150 Hz). Correlations between pairs of ROIs were used to construct functional networks.

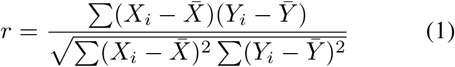

Here, *X*_*i*_ and *Y*_*i*_ represent a pictotesla or femtotesla value at index *i* measured in MEG signals of ROI x and ROI y, respectively. The Pearson correlation coefficient *r* is computed using Equation 1. It quantifies the linear relationship between the activity levels between two given regions. This is done between all regions, creating a 90 × 90 correlation matrix.

### 5.7 Identifiability metrics of individuals

Person re-identification was achieved using Equation 1, based on [4]. This is where *X* and *Y* represent the upper triangles of their functional connectomes. Each person was re-identified using correlation scores from 85 sessions. These were 2 sessions from each of the other subjects (2 × 42), and 1 session from themselves taken later. Calculations were made to evaluate intra-person identifiability (*ID*_self_), inter-person identifiability (*ID*_other_), and differential identifiability (*ID*_diff_) which was used in [26]. Inter-identifiability scores for both sessions of the same person were averaged before Equation 3. *ID*_self_ and *ID*_other_ are defined as follows:

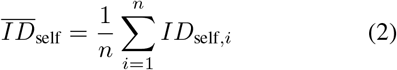

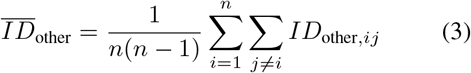

*ID*_self,*i*_ represents the average intra-identifiability measure for all subjects, and *ID*_other,*i*_ represents the average inter-identifiability measure across session scores of *j*-th other subjects.

Person recognition accuracy was evaluated and run across all subjects based on the intra-class identifiability being greater than all inter-class identifiability values. For every successful boolean comparison, a true value of 1 was taken. This is in the form of a binary recognition vector (see Equation 4, which was later summed in the fitness function:

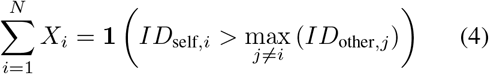

The differential identifiability is then calculated as follows:

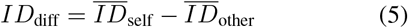

### 5.8 Optimization solution for identifying sub-networks

Simulated Annealing [40] is a probabilistic metaheuristic optimization algorithm that can approximate global optima in large search spaces via an exploratory search strategy. The algorithm was created and run with Python version 3.11.0. We modeled our optimization based on randomly swapping 10-20% regions of the total number of regions iteratively until improvement. Simulated Annealing decides the selection of a new solution based on an evaluation of fitness and probability, defined by an acceptance function.

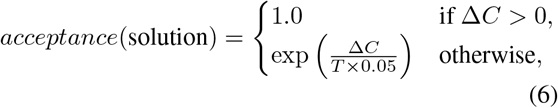

where Δ*C* = new fitness − old fitness and *T* = temperature.

The fitness function of the Simulated Annealing optimization is defined by two metrics relating to fingerprinting. Recognition accuracy, calculated from the sum of Equation 4 and differential identifiability in Equation 5 are used. In both, the minimum score of two sequential time segments of 30 s in the MEG signal is used for evaluation to ensure cross-segment consistency. A third un-optimized sequential segment of 30 s was later taken to be validated for consistency.

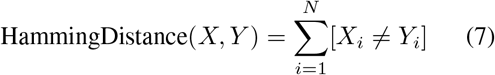

‘Hamming Distance’ (see Equation 7) is used, which measures agreement between two binary strings *X* and *Y* that represent binary recognition vectors from different segments of all 43 subjects. It is used to ensure that during convergence, the recognition outputs of a subject from different time segments are consistent.

The following is the final fitness function, which includes the following terms:

- *C*1 = binary recognitions vector of segment 1
- *C*2 = binary recognitions vector of segment 2
- *D*1 = differential identifiability of segment 1
- *D*2 = differential identifiability of segment 2
- *P* = number of persons
- HD = HammingDistance

#### Fitness maximization function

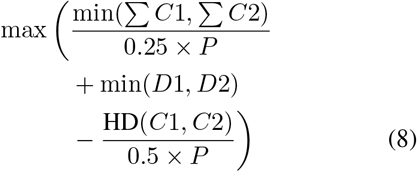

### 5.9 Graph analysis of region occurrences

Graph measures were used to observe structural patterns in the co-occurrence of regions in the optimal sub-networks of the MEG functional connectome fingerprint. The standard functions in the Python library NetworkX [41] were used to perform assortativity and clustering coefficient graph measures.

- **Assortativity** (see Methods 5.9.1) was used to observe how regions co-occur in terms of more or less region frequency occurrence.
- **Local region neighborhood clustering** (see Methods 5.9.2) showed how connected regions were to each other in a region-specific local neighborhood.
- **Region co-occurrence by subject-specific performance** (see Methods 5.9.3) showed how region co-occurrence benefited intra-identifiability scores for a subject. This showed the heterogeneity across the cohort and which regions were useful contributors to fingerprinting.

#### 5.9.1 Assortativity of connections

The assortativity tells us how regions with a lot of connections/degrees (high frequency of occurrences of regions) connect to regions of fewer connections/degrees (low frequency). Values range from −1 to 1, where - 1 shows highly connected regions connecting to lowly connected, 0 shows no relationship, and 1 is highly connected to highly connected.

The assortativity score *r* was computed using the function from the NetworkX library, corresponding to the equation used in [42]. In our approach, we calculated the degrees of a node as the sum of edge weights connected to it. In Equation 9 *x* and *y* represent the different degree scores of nodes in the graph. *E*_*xy*_ is the sum of weighted edges that connect between nodes of these degree scores. *M* represents the total sum of weighted edges in the graph. 2*M* is the total sum of weighted edge ends in the graph, which is double the number of weighted edges.

The first term 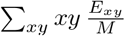 on the numerator provides a higher score when larger degrees share more summed weighted edges.

The second term 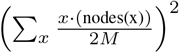 provides the expected score if the weighted edge ends from all nodes of degrees are the average quantity.

The denominator is the variance, which standardizes the value between −1 and 1.

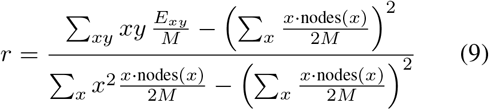

#### 5.9.2 Local region neighborhood clustering

The clustering coefficient tells us how connected one region and its local neighborhood are. It shows whether regions co-occurring in the region of interest also co-occur with each other.

The local clustering coefficient was computed using the NetworkX library. The local clustering coefficient *c*_*u*_ for a region *u* is given where *T* (*u*) is the number of triangles where two other connected nodes with *u* also connect with each other. The denominator shows the maximum number of triangles through *u*, where *deg*(*u*) is the degree of the node *u*:

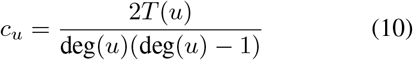

#### 5.9.3 Region co-occurrence by subject-specific performance

The intra-identifiability between all regions was taken, and for each region, the maximum value of its edges was taken to be its best-performing co-occurrence.

Degree divergence and leaf number were then taken from the graph constructions to show the distribution of optimal co-occurring regions.

The degree divergence represents the broadness of degree distribution, used in [43]. This represents the broadness for the same regions to be optimal in co-occurrence in each subject. Degrees are how many edges a node/region is connected, where ⟨ ⟩ indicates the average:

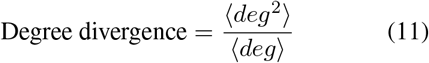

The leaf number represents the number of regions that did not maximally co-occur with any other regions. This value is taken out of 89.

### 5.10 Machine learning model for disease differentiation

Our deep learning model was trained using TensorFlow version 2.16.1. To perform disease differentiation, sub-networks identified from the Simulated Annealing optimization on the CUBRIC dataset were re-applied to the OMEGA dataset, obtaining 46,071 10 × 10 correlation matrices for all 30 Parkinson’s patients and 30 healthy subjects. After taking only the upper triangle from the correlation matrix, each sub-network represented a vector of length 45 values.

The 60 individuals provided 46,071 sub-networks each. The disease differentiation patterns were learned from binary classification on a dual-objective stacked auto-encoder model architecture (see Supplementary Methods 2). The model was trained for 100 epochs for each sub-network feature selection and each test fold. 6-fold cross-validation was used across individuals where the sub-networks of 5 Parkinson’s and 5 healthy subjects were treated as untouched testing samples. F1 Score is the harmonic mean of precision and recall metrics (see Supplementary Methods 3), which allows us to obtain a single metric for the differentiation of diseased from healthy.

The dual-objective stacked autoencoder was trained on 40,000 of the 46,071 optimal fingerprint sub-networks from this study and 40,000 sub-networks from a null distribution of randomly selected regions.

## Data availability

Data for the region indices in all 46,071 optimal sub-networks and 24,197 independently sampled sub-networks from the AAL atlas that were generated and analyzed in this study will be made available in the published paper.

## Code availability

All codes for the Simulated Annealing optimization and deep learning model are available on https://github.com/VasilesBalabanis/MEGSubnetworks and Zenodo repository.

## Acknowledgements

The UKRI Centre for Doctoral Training in Artificial Intelligence, Machine Learning & Advanced Computing (AIMLAC) supports and funds this work.

We acknowledge the support of the Supercomputing Wales project, which is part-funded by the European Regional Development Fund (ERDF) via the Welsh Government.

We acknowledge the help of Dr Michael Kenning for graph-related discussions on the investigation of sub-networks.

## Author contributions

All authors conceptualized the study. V.Balabanis performed the analyses and wrote the first draft of the manuscript. All authors contributed to the writing and editing of the manuscript.

