## Supplementary Materials for "Robust sub-network fingerprints of brief signals in the MEG functional connectome for single-patient classification"

### 1. Participants' Summary

Supplementary Table 1: **Summary of CUBRIC Multi-Session Dataset**. Additional details of the MEG sessions provided by Cardiff University.

| Characteristic | Value |
|---|---|
| Total number of subjects | 43 healthy subjects |
| Sex | 12 men, 31 women |
| Duration between Sessions | Mean: 6.07 days, ± 6.14SD, Range: 1-27 days |
| Age (for 37 subjects) | Mean: 22.45, ± 3.06SD, Range: 18-30 |
| Experiment | 5 minutes resting-state per session |

Supplementary Table 2: **Summary of OMEGA Single-Session Dataset**. Details of the open-access MEG sessions provided by the Open MEG archive.

| Characteristic | Value |
|---|---|
| Total number of subjects | 60 (30 patients, 30 healthy subjects) |
| Healthy subjects' Age | Mean: 67.11, ± 8.35SD, Range: 50.52-82.85 |
| Parkinson's patients' Age | Mean: 67.35, ± 8.88SD, Range: 48.53-80.14 |
| Experiment | 5 minutes resting-state |

### 2. Neural Network Architecture

The following deep learning architecture was used to learn the patterns in the sub-networks in the MEG functional connectome of the brain. The activation functions used were Rectified Linear Units (ReLU) [44], known for their preservation of gradient values. Drop-out (de-activation of neurons), batch-normalization [45], and L2 regularization were used to prevent the model from overfitting.

#### 2.1. Encoder

Supplementary Table 3: **Encoder layer of the deep learning model.** This is the backbone layer which provides input to the decoder and classifier.

| Layer | Neurons | Equation |
|---|---|---|
| Input | - | $\mathbf{x} \in \mathbb{R}^{45}$ |
| Dense | 512 | $\mathbf{z}_1 = \text{ReLU}(\mathbf{W}_1\mathbf{x} + \mathbf{b}_1)$ |
| Batch Norm & Dropout | - | $\mathbf{z}'_1 = \text{Dropout}(0.3)(\text{BatchNorm}(\mathbf{z}_1))$ |
| Dense (L2 reg.) | 256 | $\mathbf{z}_2 = \text{ReLU}(\mathbf{W}_2\mathbf{z}'_1 + \mathbf{b}_2 + \lambda\|\mathbf{W}_3\|^2)$ |

#### 2.2. Decoder

Supplementary Table 4: **Decoder layer of the deep learning model.** This is the output layer which reconstructs the input

| Layer | Neurons | Equation |
|---|---|---|
| Dense | 256 | $\mathbf{z}_3 = \text{ReLU}(\mathbf{W}_3\mathbf{z}_2 + \mathbf{b}_3)$ |
| Batch Norm & Dropout | - | $\mathbf{z}'_3 = \text{Dropout}(0.3)(\text{BatchNorm}(\mathbf{z}_3))$ |
| Output | 45 | $\hat{\mathbf{x}} = \sigma(\mathbf{W}_4\mathbf{z}'_3 + \mathbf{b}_4)$ |

#### 2.3. Classifier

Supplementary Table 5: **Classifier layer of the deep learning model.** This is the output layer which binary classifies a sub-network as diseased or healthy.

| Layer | Neurons | Equation |
|---|---|---|
| Dense | 128 | $\mathbf{y}_1 = \text{ReLU}(\mathbf{W}_5\mathbf{z}_2 + \mathbf{b}_5)$ |
| Dropout | - | $\mathbf{y}'_1 = \text{Dropout}(0.3)(\mathbf{y}_1)$ |
| Output | 1 | $\hat{\mathbf{y}} = \sigma(\mathbf{W}_6\mathbf{y}'_1 + \mathbf{b}_6)$ |

#### 2.4. Loss and Optimization

Mean Squared Error (MSE) loss was used across each batch of N samples of the 45-dimensional input vector:

$$\text{MSE} = \frac{1}{N}\sum_{i=1}^{N}\frac{1}{45}\sum_{j=1}^{45}(\hat{x}_{ij} - x_{ij})^2$$

Binary Cross-Entropy (BCE) loss was used for a 1-D output across each batch of N samples:

$$\text{BCE} = -\frac{1}{N}\sum_{i=1}^{N}[y_i \log(\hat{y}_i) + (1 - y_i)\log(1 - \hat{y}_i)]$$

$$\text{Cost function} = BCE + MSE$$

The Adam (adaptive moment estimation) optimizer [46] was used for gradient descent based on its known robust and adaptive approach to learning rate adjustments.

### 3. Evaluation Metrics

The evaluation metrics for the binary classification (Positives referring to diseased and Negatives referring to healthy) are shown below.

Accuracy is the ratio of correct classifications (both diseased and healthy) from all subjects/patients:

$$\text{Accuracy} = \frac{\text{True Positives} + \text{True Negatives}}{\text{Total Observations}}$$

Precision is the ratio of correctly classified patients to all subjects/patients classified with the disease:

$$\text{Precision} = \frac{\text{True Positives}}{\text{True Positives} + \text{False Positives}}$$

Recall is the ratio of correctly classified patients to all actual patients:

$$\text{Recall} = \frac{\text{True Positives}}{\text{True Positives} + \text{False Negatives}}$$

F1 Score is the harmonic mean of precision and recall, which allows us to obtain a single metric for the differentiability of diseased from healthy:

$$\text{F1} = 2 \times \frac{\text{Precision} \times \text{Recall}}{\text{Precision} + \text{Recall}}$$

#### 4. Supplementary Figures

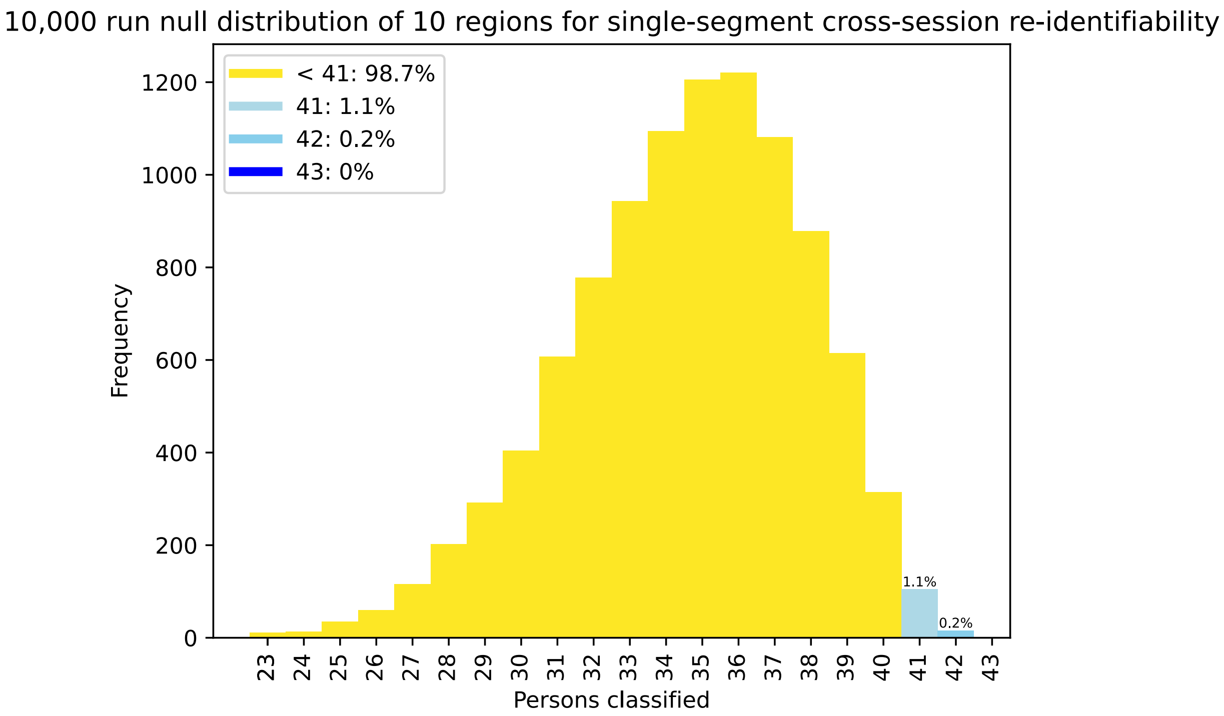


Supplementary Figure 1: **Null distribution of 10,000 runs on 10 regions.** This shows the performances in 41/43+ classifications (95% accuracy) composing less than 5% of the overall distribution.

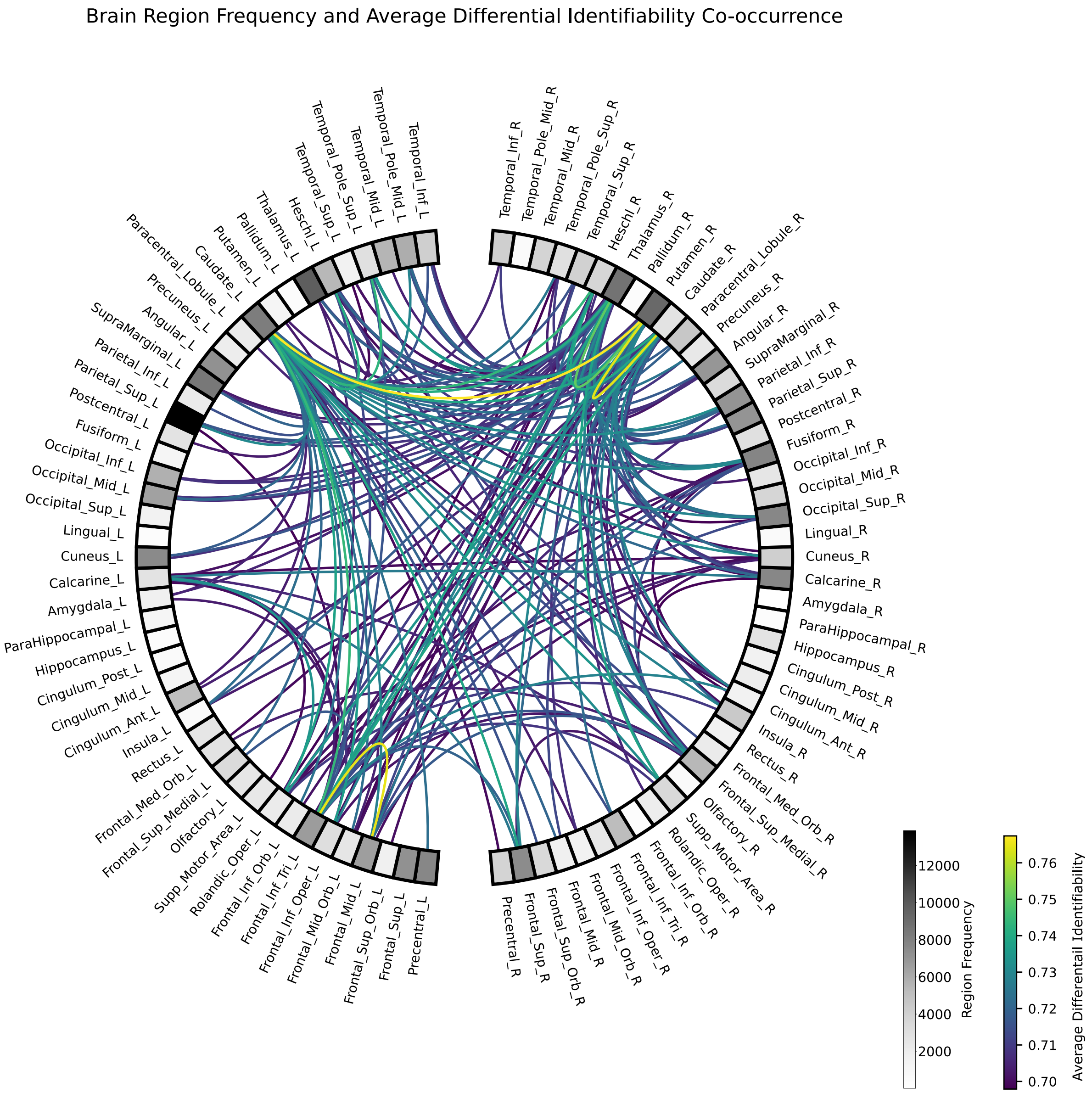


Supplementary Figure 2: **Average co-occurrence differential identifiability.** Values are thresholded to the top 10% of differential identifiability scores, including only where the co-occurrence frequency is at least 100 for confidence.

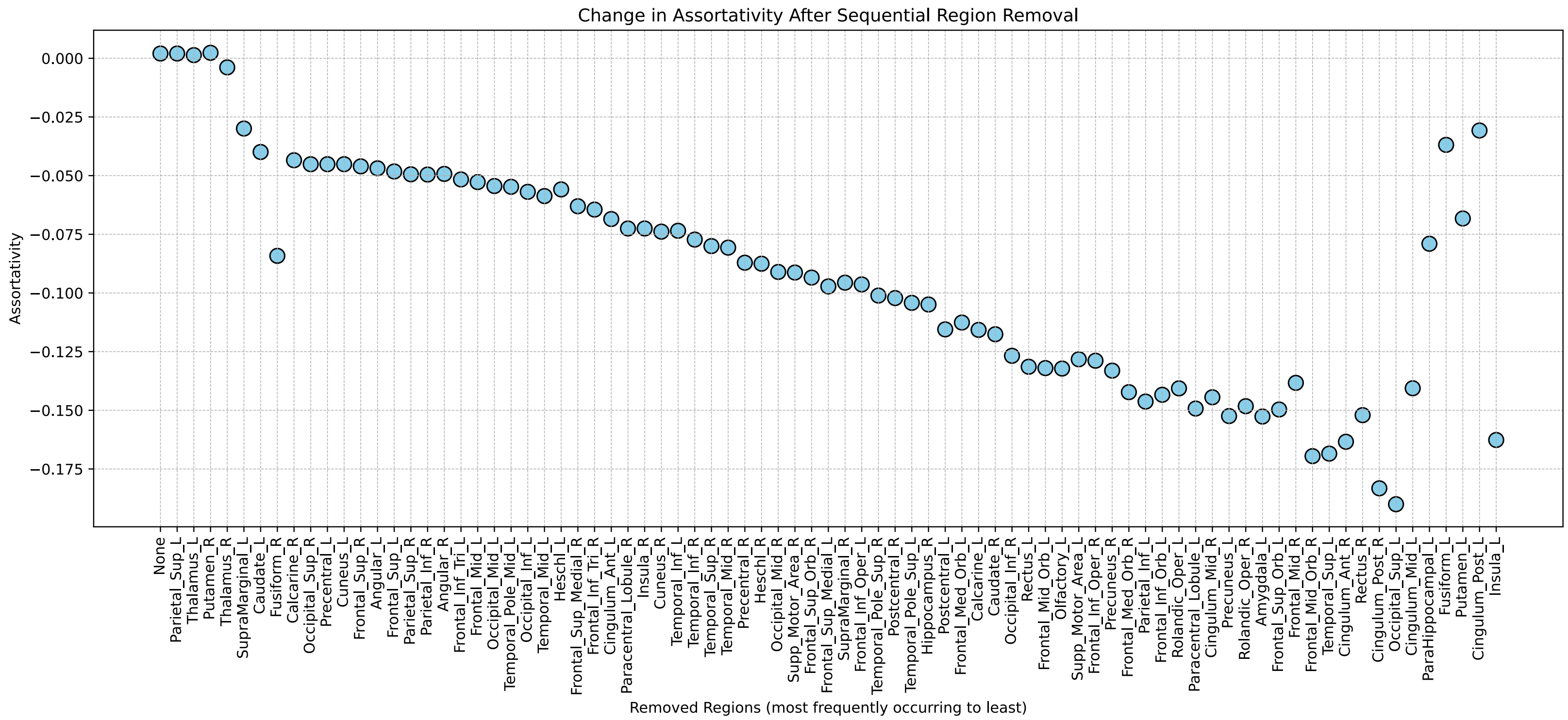


Supplementary Figure 3: **Assortativity of all 24,197 sub-networks constructed as a full graph** (see Methods 5.9.1). Sequential region removal is done based on the most frequently occurring regions. This is to observe an expected maximal fluctuation in assortativity.

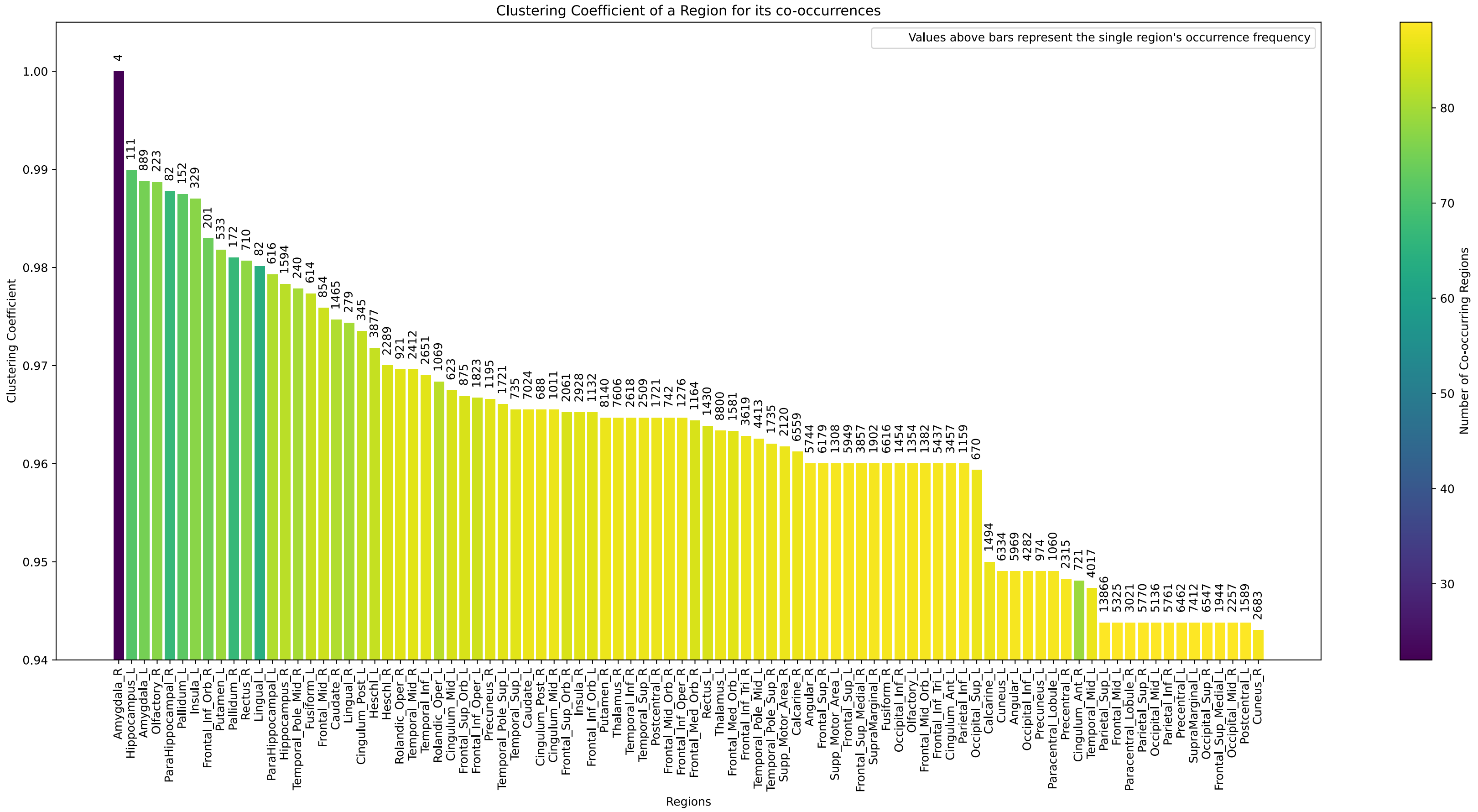


Supplementary Figure 4: **The clustering of a region's neighborhood with itself** (see Methods 5.9.2) on the 24,197 sub-networks in our results. Regions may also not co-occur with all regions, shown as a darker hue.

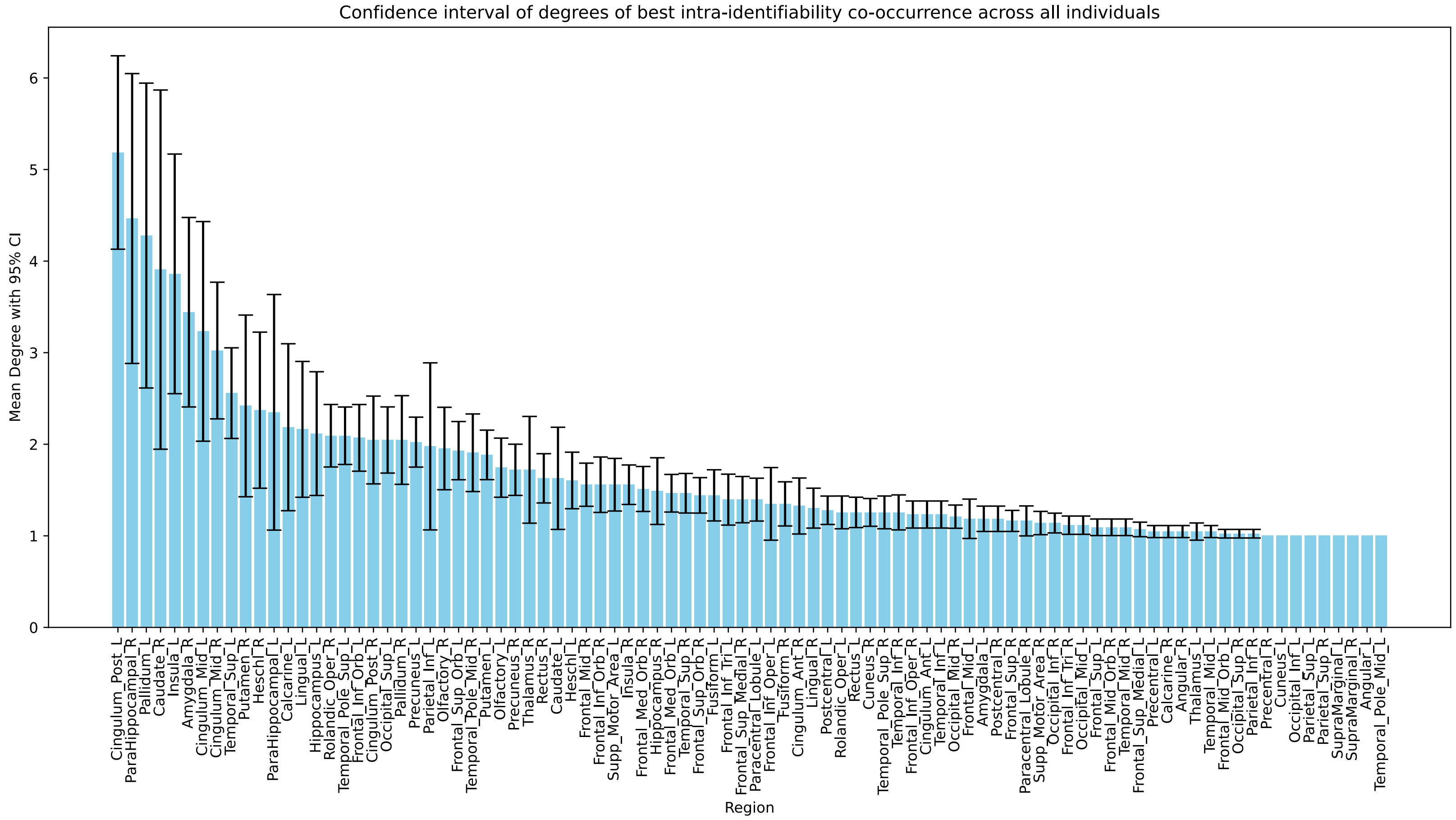


Supplementary Figure 5: **Degree distribution across 43 subjects in a graph construction** (see Methods 5.9.3). Degrees are based on the number of edges of a particular region, which are cumulatively added when the averaged intra-identifiability of the region co-occurrence is higher than all other regions. This was done on 24,197 independently sampled sub-networks.

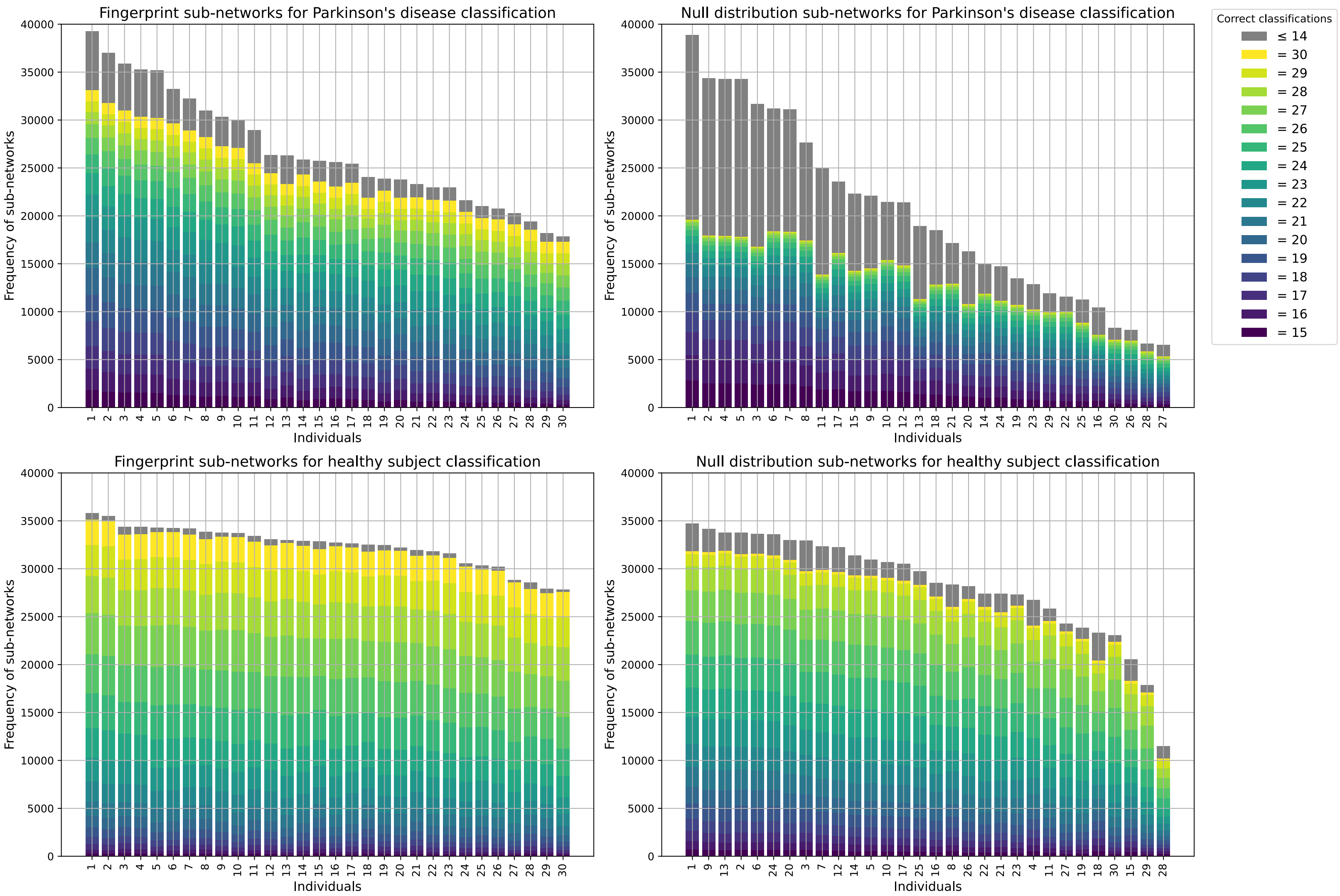


Supplementary Figure 6: **Binary test classifications of Parkinson's disease sub-networks and healthy sub-networks using the dual-objective stacked autoencoder** (see Methods 5.10). The number of successful classifications of each individual is shown as the score of each bar. The colored sections of each bar represent the consistency of the sub-networks for classifying all individuals in the group.
